# A system for 3D reconstruction of mouse behaviour

**DOI:** 10.1101/2025.11.05.686730

**Authors:** Storchi Riccardo

## Abstract

Recent development in machine vision has enabled researchers unprecedented resolution in measuring behaviour in freely moving animals. However, the hardware implementation of a behavioural arena for capturing 3D behaviours remains time consuming and unstandardised. Here we first provide detailed instructions on how to build and calibrate such a system. Secondly, we provide instructions on how to process video data to obtain a 3D reconstruction of mouse behaviour. Finally, we provide results from behavioural tests in which 3D reconstruction enables clear-cut separation of behavioural responses to distinct sensory stimuli.

**Data and Software Availability:** All data and analyses software required for this article is provided in a publicly available repository [1]

**Competing Interests:** This project contains the following underlying data.

**Grant Information:** This work was funded by NC3Rs via a David Sainsbury Fellowship (Project Reference NC/P001505/1).

## Introduction

The advent of deep learning and powerful unsupervised classification algorithms has enabled researchers to automatically track multiple body landmarks in freely moving animals [2, 3] and generate detailed and unbiased behavioural classifications [4-10]. This has provided three fundamental opportunities.

Firstly, the capacity to quantify and understand complex behaviours. Thus, by reliably tracking multiple parts of the body and classifying posture and movements, we can start to understand not only where an animal is but also what an animal is doing and its internal state. Over the last few years these techniques have found many applications in rodent studies empowering researchers to capture arousal and emotional states [11, 12], age and sex differences [13], body postures and movements [5, 6, 8, 10, 14, 15], eye and whisker movements [16, 17], defensive [5] hunting [18-20] and social behaviours [21, 22].

Secondly, the possibility to reduce the number of animals required for testing specific treatments, or genotypes. Automatic and detailed classification of mouse behaviour substantially increases the ability to discriminating the effects of pharmacological treatments compared with standard metrics [7]. Similarly, it increases the ability to discriminate mouse genotypes [8, 9, 23, 24] and provides better discrimination of specific avoidance behaviours [5]. This has clear advantages for reducing animal use.

Thirdly, the increased throughput and accuracy in quantifying natural freely moving behaviours suggests the possibility to replace, at least in part, head-fixed experiments. This is a significant refinement for animal welfare since head-fixation can be stressful and requires habituation sessions [25]. Moreover, freely moving experiments often rely on ethologically relevant innate behaviours (e.g. exploration, avoidance, hunting) and therefore don’t require training procedures involving food and/or water deprivation.

Quantification of behaviour is typically performed with a single camera mounted on top of a behavioural arena. Since cameras project a 3D space onto a 2D image the resulting videos cannot capture physical distances and crucially depend on the positioning of the camera. Thus, while sufficient to capture a rich behavioural repertoire, this approach cannot accurately quantify body coordinates, head and body postures, or vertical movements (e.g. rearing, sniffing).

The solution to these problems is to employ multiple cameras to obtain a 3D reconstruction of the animal and its surrounding environment. Yet, building a multicamera system and calculating 3D coordinates introduces further complications such as building a multicamera frame, synchronising the cameras, calibrating a multicamera system, correcting for outliers in 3D reconstruction. In this article we provide a detailed list of instructions to overcome these problems. At the end of each step required for setting the system, we provide the list of components required, sample codes and links to the relevant code repositories.

## Materials and Methods

### Step 1 - Building a 3D frame

Building a stable frame is fundamental for all the following steps. Schematics and pictures of the 3D frame are shown in **Figure 1**. To do that we used 25 mm optical rails that can be easily assembled with a standard Allen Keys set. To connect the rails, we used angled brackets and counterbored construction cubes. Both types of connections rely on low profile channel screws and drop-in T-nuts (see **Table 1**).

**Table 1:**
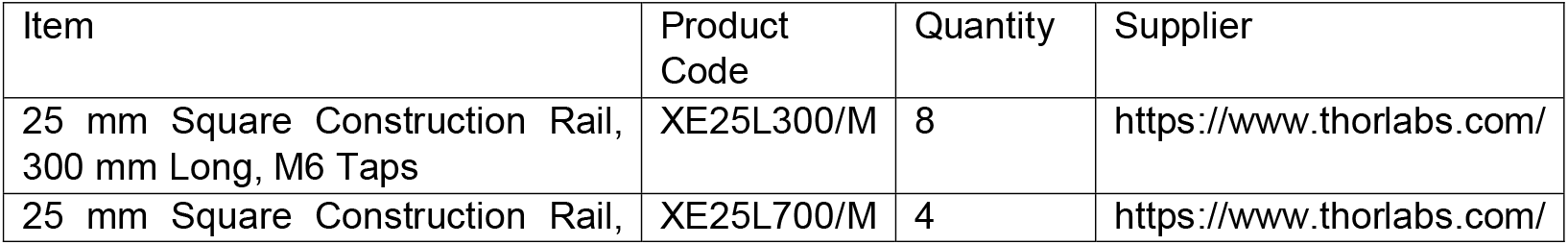
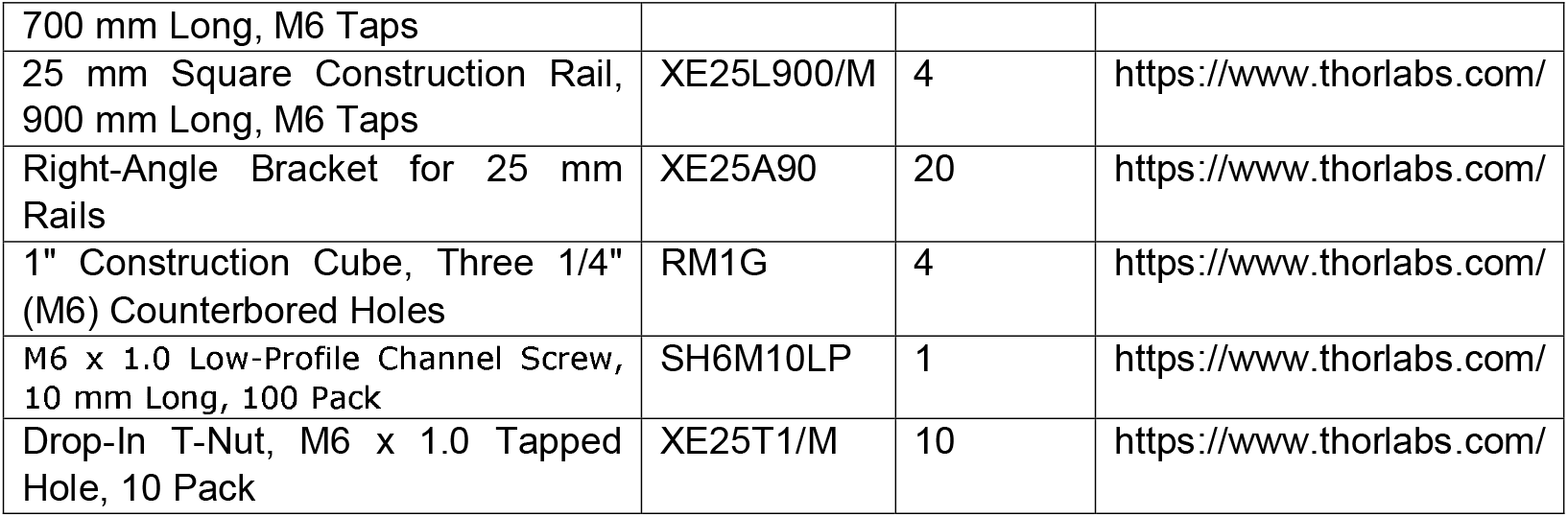
List of components required for building the 3D frame.

**Figure 1:**
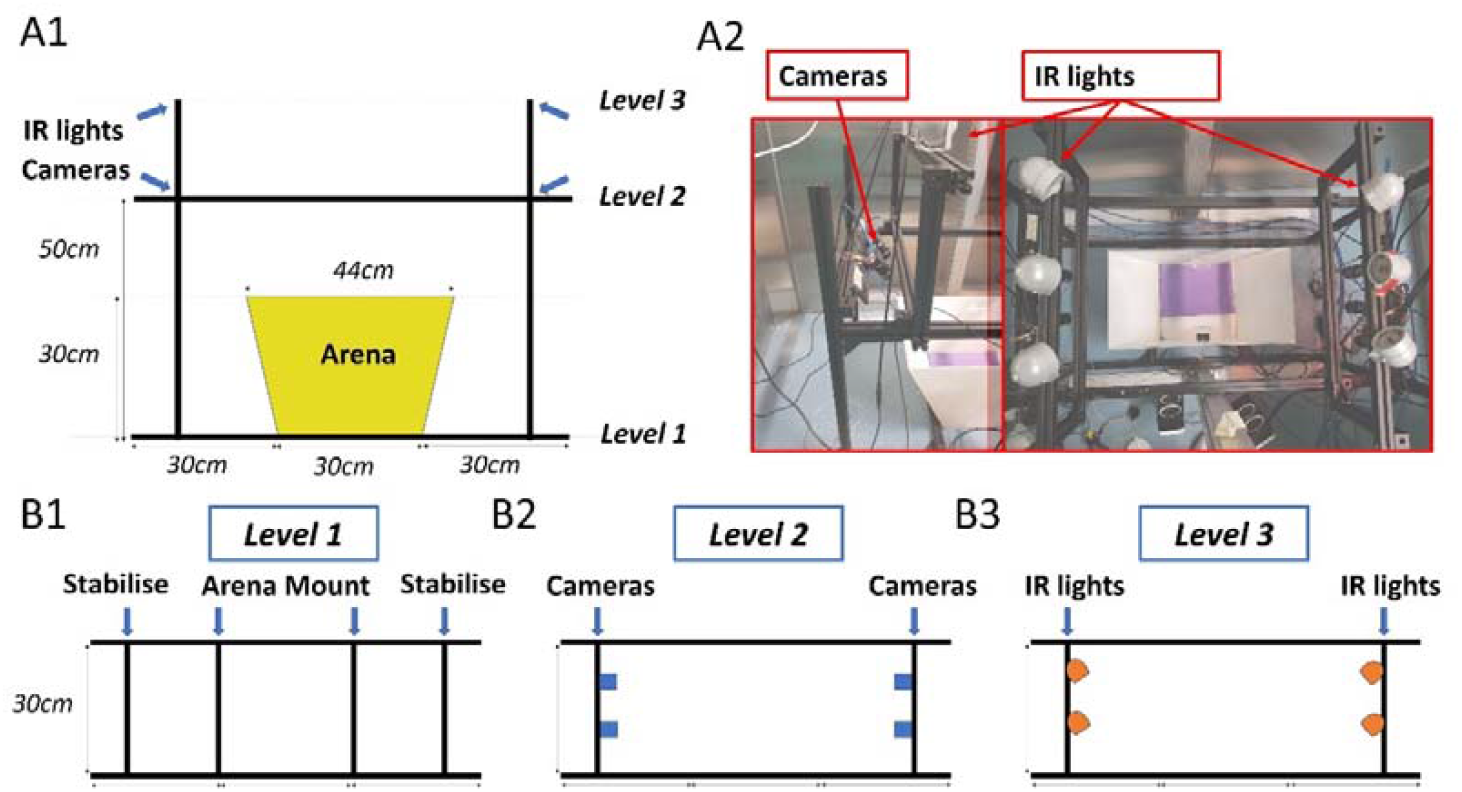
A1) Schematic of the 3D frame (side view). A2) Side (left panel) and top view (right panel) of the 3D frame. B1-3) Schematic of the 3 different levels of the 3D frame (top views).

A basic frame for a 30cm arena requires the following elements: 4 horizontal long rails, 4 vertical long rails, 8 horizontal short rails. A drawing of the frame containing a 30cm-base behavioural arena is shown in **Figure 1A1** from a side view. There are 3 vertical levels (see ***Level 1***,***2***,***3*** in **Figure 1A1**). At the ***Level 1*** we use two short rails to stabilize the structure and two to keep the arena in the specified position (**Figure B1**). At the ***Level 2*** we use two short rails to mount the cameras on (**Figure B2**). Each short rail mounts two cameras (see “Assembling the cameras” section). At ***Level 3*** we use two short rails to mount the infrared lights. A picture of the full system is shown in (**Figure B3**).

### Step 2 - Building the arena

The shape of the arena has a significant effect on the quality of video recordings. Occlusions (blind spots) should be avoided to make sure that each camera can capture the animal in all locations of the arena. To achieve that we built a “boat-shaped” arena, with the top horizontal dimensions larger than the bottom one (**Figure 1A**). In this way it is possible to avoid blind spots typically occurring when animals seek shelter near the walls or the arena corners. The easiest way to build the arena is to order 5 rectangular panels of Perspex (thickness = 5mm) and cut the two trapezoidal ones in a workshop (dimensions shown in **Figure 1A1**). Tracking works best when floor and wall reflections are minimised. To do that we sanded the internal walls of the arena.

### Step 3 - Assembling the cameras

We used four cameras for computer vision. A full list of the accessories required, including lenses and filters, is provide in **Table 2**. We applied C-mount fixed focal lenses. Since the cameras are CS mount, we a C-mount to CS-mount adapter to focus the images on the camera sensor. Variable conditions of illuminations can make tracking harder, or impossible when camera sensors saturate. To avoid this problem, we supplied the arena with constant infrared illumination (see **IR Lights** in **Figure 1**) and a high-pass optical filters to prevent visible light to reach the camera sensors.

**Table 2:**
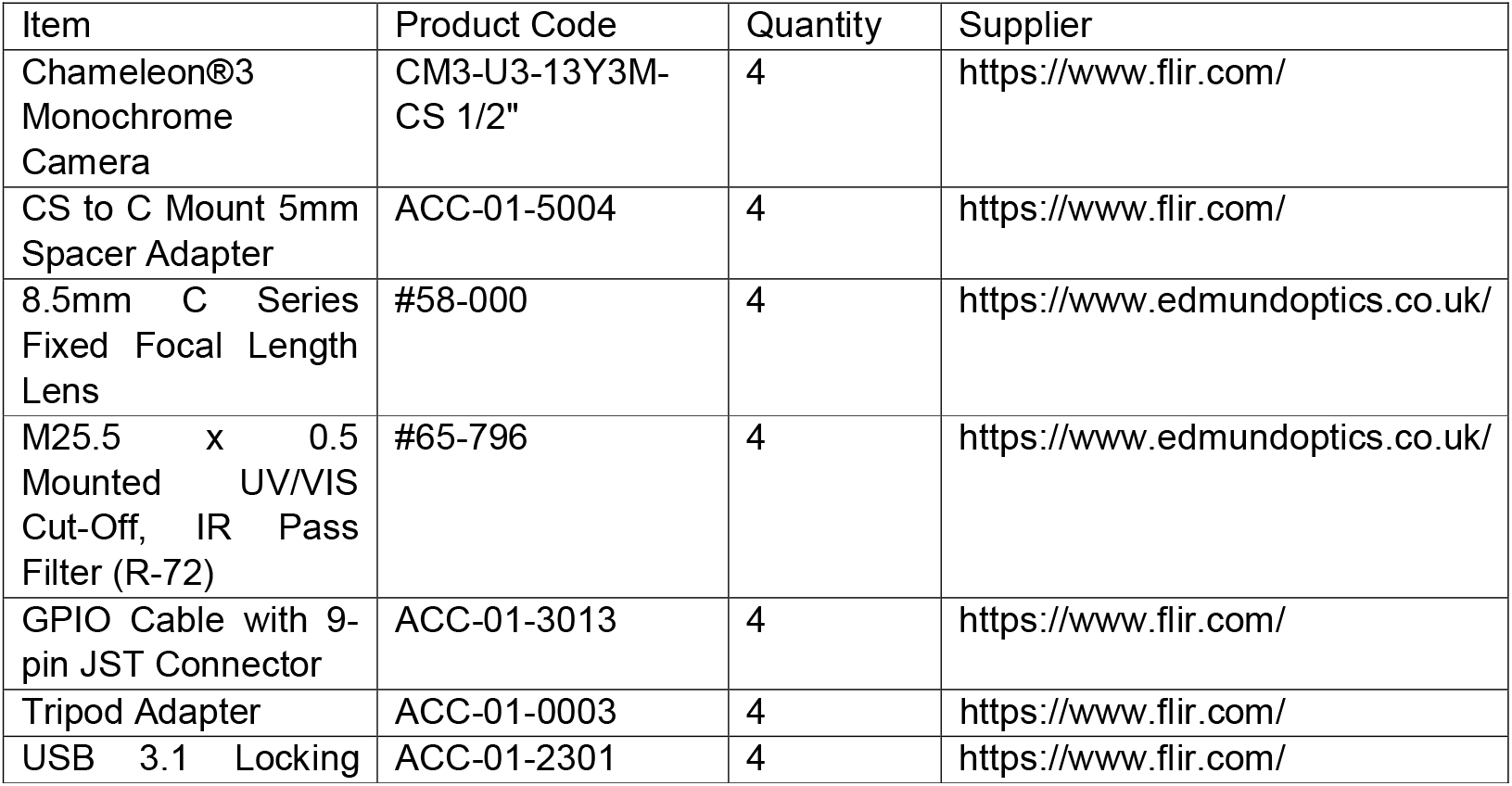
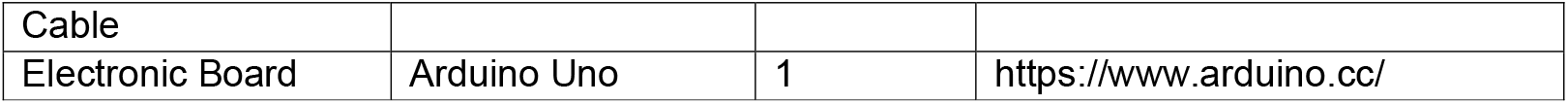
List of components required for the four cameras.

### Step 4 - Synchronising the cameras

To accurately triangulate body landmark coordinates detected by different cameras it is essential that camera frames are tightly synchronised. To achieve that we used an electronic board (Arduino Uno, see **Table 2**) to deliver single TTL pulses to all cameras. To connect the board to each camera we used JST connectors (see **Table 2**) and soldered yellow and brown wire respectively to a digital input (we used digital input 3, see **Box 1**) and a ground of the electronic board. A sample code for the electronic board is provided in **Box 1**. To upload the code on the board, Arduino software (here we used version 1.8; https://www.arduino.cc/; newer versions also work) must be installed and the board connected to a PC via a USB cable. Uploading only needs to be performed once. After that, the board will run the same code every time is turned on.

#### Box 1: Sample code to upload to the electronic board

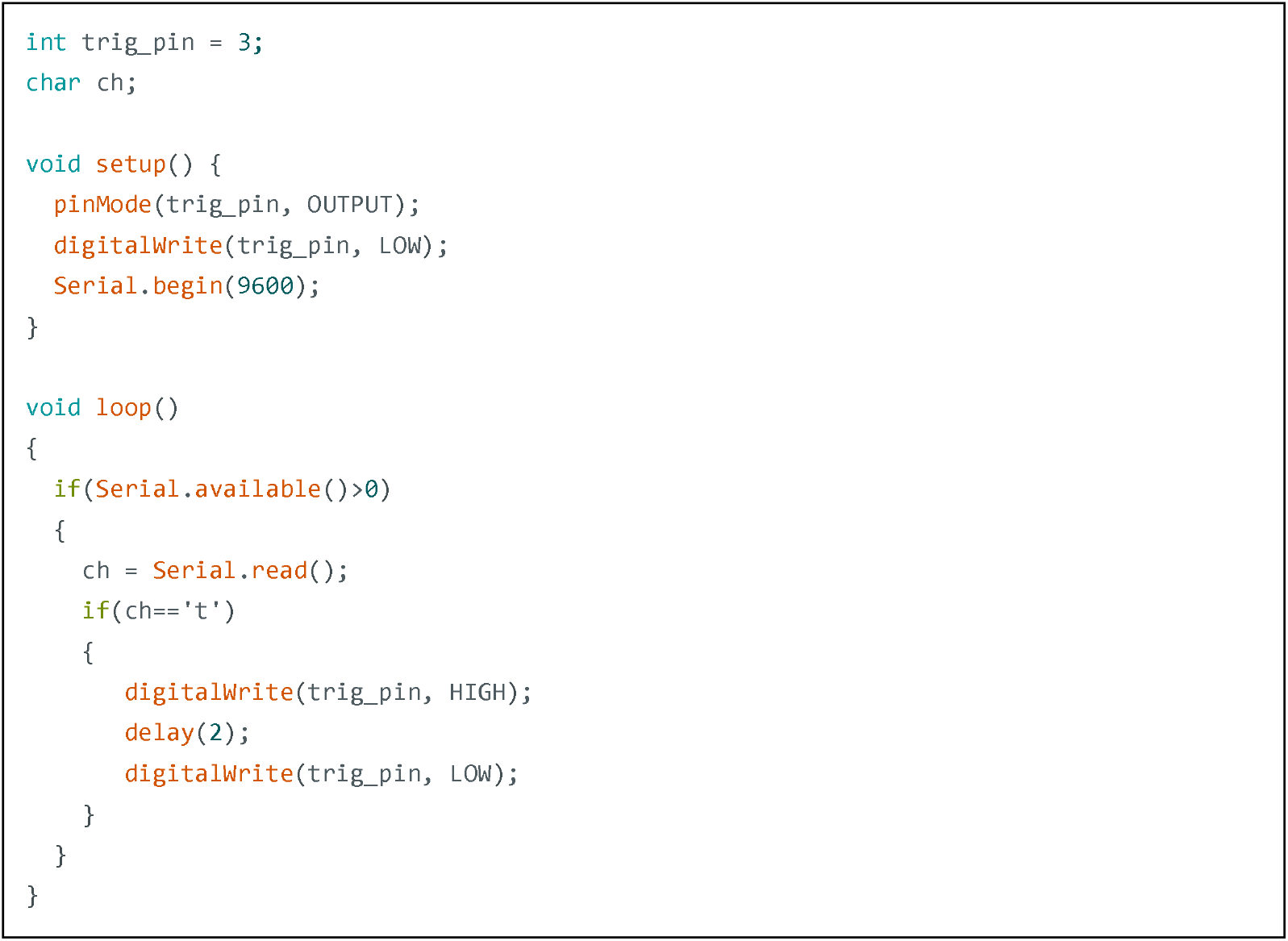

### Running a multicamera acquisition

Install **Point Gray FlyCap2** software (here we used version 2.13.3.61; newer versions also work). This will automatically recognise the cameras once those are connected to the PC. Make sure cameras are connected to USB3 ports as USB2 ports will substantially reduce the maximum frame rate of your acquisition. If the acquisition is performed with a laptop lacking 4 USB3 ports you can use a USB 3.1 Hub (Product Code: ACC-01-6006 from https://www.flir.com/). Open four instances of the program, one per each camera. Under *Settings* select the preferred format. Under *Settings* tick the box “*Enable TTL*”. This will stop video acquisition until TTL triggers are provided. Under *Capture* select *Video*. Leave frame at 0.

To run a multicamera acquisition, any programming software that can write to a serial port would work. Here we report sample code for Python (**Box 2**). To successfully run the code, take a note of the serial port number (e.g. *‘COM3’* in our example, see **Box 2**) which can be found with PC’s *Task Manager*. Edit the sample code to match this number.

#### Box 2: Sample code to run a multicamera acquisition

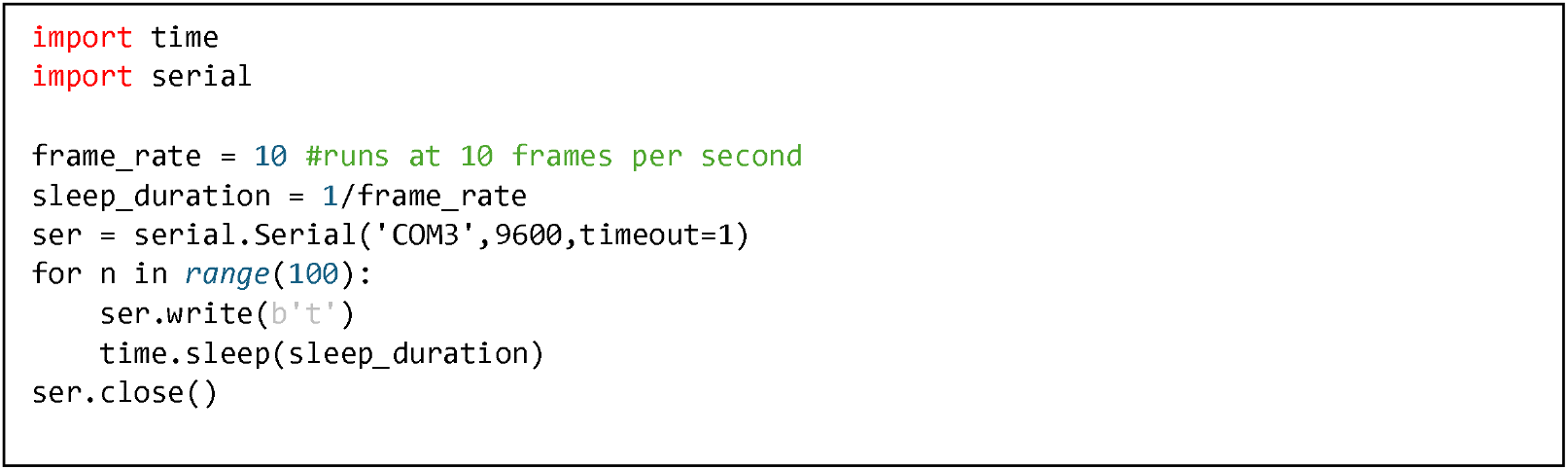

### Step 5 - Calibrating the multicamera system

Several methods have been developed to calibrate multicamera systems. A classical approach is to employ a printed checkboard and toolboxes to perform these calibrations are available in OpenCV and MATLAB. Here we preferred to use 3D physical objects of known size (LEGO®) and custom written MATLAB code. The scripts required to perform the calibration routine are provided at this link [1] within the *3D calibration* folder.

To run the calibration first prepare the LEGO® plates. Use dark plates and bricks and mark the relevant coordinates with a black marker pen to increase their visibility (see **Figure 2**). We used a total of five plates, one with no bricks the others with bricks of different heights (**Figure 2**). Take a picture (in *.tif format) of each plate with each camera. Make sure each plate is placed at the same exact location as doing otherwise will affect the calibration results.

**Figure 2:**
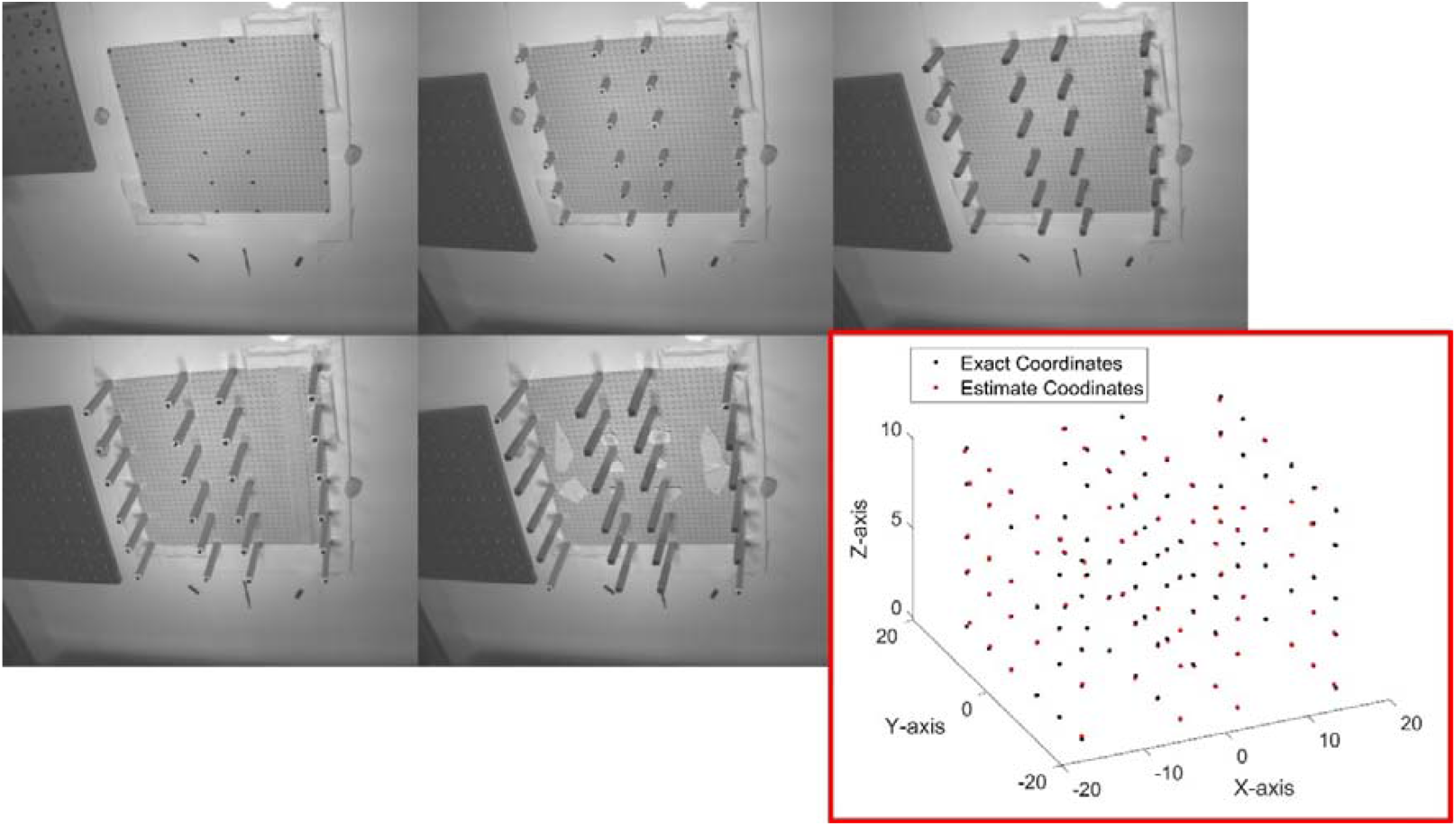
Pictures of the 5 LEGO® plates taken from one of the four cameras. Bottom right panel shows the 3D reconstruction of the LEGO® coordinates after calibration (exact and estimated coordinates shown in black and red dots).

The code to run the calibration routine is provided in [1] and to run it follow instructions provided in *CalibrationInstructions*.*ppt*. Once calibration is finished you can estimate the camera matrices, required for 3D reconstruction, with the script *calibrate_cameras_cl*, also provided in the same repository. The script will save a file (*Pcal*.*mat*) with camera matrices and also output a figure comparing real world and 3D reconstructed coordinates (**Figure 2**, bottom right panel). A good match between these sets of coordinates is required to obtain reliable 3D reconstructions.

| Item | Product Code | Quantity | Web Link |
| --- | --- | --- | --- |
| Plate | BASE PLATE<br>32X32 | 5 | https://www.lego.com/en-gb/pick-and-build/pick-a-brick |
| Bricks Height 3 | BRICK 1X1X3 | 48 | https://www.lego.com/en-gb/pick-and-build/pick-a-brick |
| Bricks Height 5 | BRICK 1X1X5 | 72 | https://www.lego.com/en-gb/pick-and-build/pick-a-brick |

### Step 6: Initial 3D reconstruction

In our set-up performing a 3D reconstruction required 2D tracking data from four cameras. Use DeepLabCut (here we used version 1; newer versions also work) [26] to perform the tracking of videos from individual cameras. This software will provide four files in csv format. Edit the script *triangulate_tracked_data*.*m* in [1] repository, *3Dcalibration* folder) to provide the names and path of these files (lines 9-12) and add the name and path of the calibration file obtained under **Step 5** (line 15). Running the script will return a 3D reconstruction of the animal body coordinates.

### Step 7: Refinement of the 3D reconstruction

The initial 3D reconstructions obtained under **Step 6** can be contaminated by outliers (see a discussion of this problem in [27]). Appropriate placement of recording cameras is important to minimise this problem. Additionally, the system we describe is scalable more cameras can be used to minimise outliers with minimal modifications. Several algorithms have been developed to identify and correct outliers and the code has been made available in public repositories [4, 6, 27, 28].

### Behavioural Experiments

Data were collected from C57BL/6 adult male mice (n = 29) provided by the Biological Service Facility at the University of Manchester. Animals were kept on a 12:12 light/dark cycle, housed in cages of 3 and provided with food and water *ad libitum*.

A detailed description of all relevant parameters to replicate the behavioural experiments is provided in [5]. Animals were initially transferred to the arena described in this manuscript by tube handling [29] and were allowed 10 minutes of habituation to the novel environment. Then, visual (looming or a flash) or auditory stimuli (a white noise or a pure tone) were delivered with an interstimulus interval of 70s. Each stimulus duration was set at 1s. The order of stimuli presentation (n = 18 stimuli per animal) was block randomised and each block was constituted by one flash, one looming and one auditory stimulus. Running of the experiment and generation of the stimuli was controlled by Psychopy (here we used version 1.82.01; https://www.psychopy.org/; newer versions also work).

All videos were acquired by 4 synchronised cameras (as described above) at 15 frames/s. Tracking of animal body landmarks (n = 5; nose, two ears, neck base and tail base) was performed with DeepLabCut [26] (version 1) and 3D reconstruction by custom-made software based on 3D Statistical Shape Models [5]. Since all analyses were automated no blinding strategy was implemented.

## Results

Here we used the 3D reconstructed data to visualise and compare spontaneous behaviours and behavioural responses to two different visual stimuli (a flash and a looming disc) and an auditory stimulus. Stimuli duration was 1 second. An in-depth description of the stimuli and all 3D data are provided in [5] and in the repository [1].

Briefly, behavioural features were extracted by fitting a 3D statistical shape model to quantify locomotion speed, body shape and orientation (see [5] for details). This analysis returned separate time series for body rearing, for the first two body shape parameters (body elongation and body left/right bending) and for locomotion speed. The time series were then segmented into behavioural epochs (2 seconds per epoch, 30 samples per each behavioural feature). An illustrative example of a single behavioural epoch is shown in **Figure 3A**. Thus, by combining rearing, shape parameters and locomotion each behavioural epoch was represented by 120 dimensions. Across our dataset of 29 animals, we collected 1032 behavioural epochs during different behavioural states encompassing spontaneous exploration, visual- and auditory-evoked behavioural responses. The dimensionality of behavioural epochs was first reduced by applying a principal components analysis that captured >90% variance within the first 20 components (**Figure 3B**). We then the applied the t-distributed Stochastic Neighbour Embedding algorithm (t-SNE, [30]) to embed these 20 dimensions in a two-dimensional t-SNE space (**Figure 3C**).

**Figure 3:**
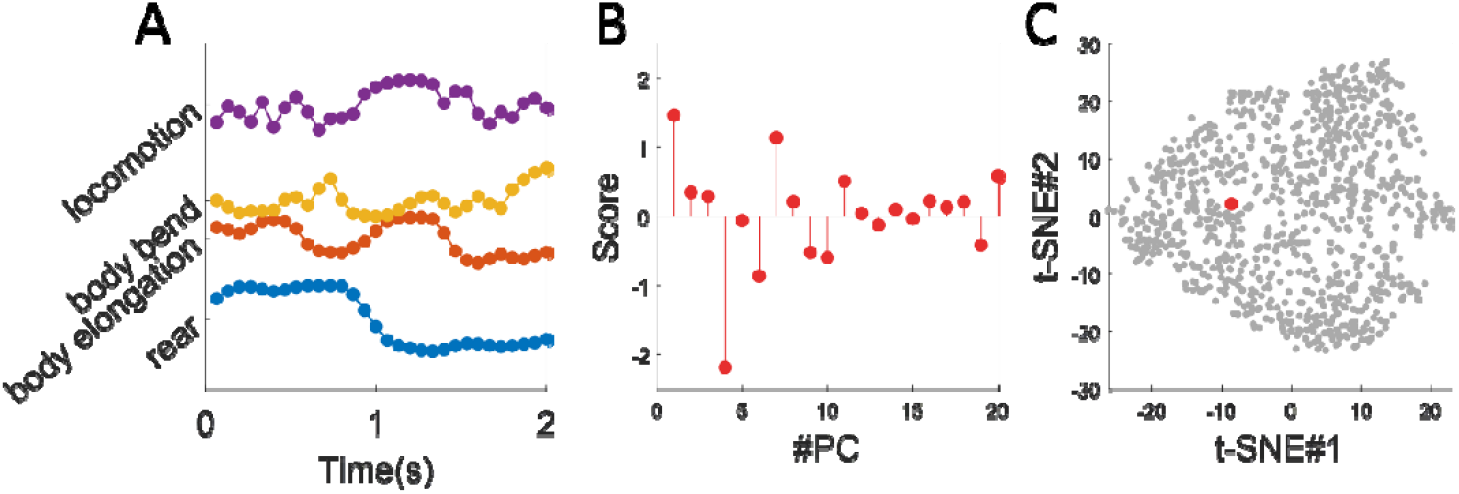
**A**) A single behavioural epochs (n = 120 dimensions) encompassing time series for locomotion speed, body shape (body elongation; body left/right bend) and rearing. **B**) The dimensionality of behavioural epochs was first reduced by applying a principal component analysis (PCA). Here we show the first 20 PCA scores for the epoch in panel **A. C**) PCA scores were embedded in a 2D space by applying the t-SNE algorithm. Here we show the embedding of the epoch in panel **A** (red dot) and for all the other epochs in our dataset (grey dots).

These preliminary steps (principal components followed by t-SNE embedding) enabled to map the high, 120-dimensional behavioural epochs (**Figure 3A**) into a 2D representation (**Figure 3C**). To compare across different behavioural states, we used the points in the t-SNE space to generate 2D count histograms, hereafter called t-SNE maps (**Figure 4A**). Visual inspection of the t-SNE maps suggests that sensory evoked responses (Flash, Looming, Sound; see **Figure 4A**) are segregated on different regions of the map, albeit with some overlap. It also shows that spontaneous exploration (Spontaneous, see **Figure 3A**) spans a wider range of behaviours that partially overlap with the more constrained set of sensory evoked responses.

**Figure 4:**
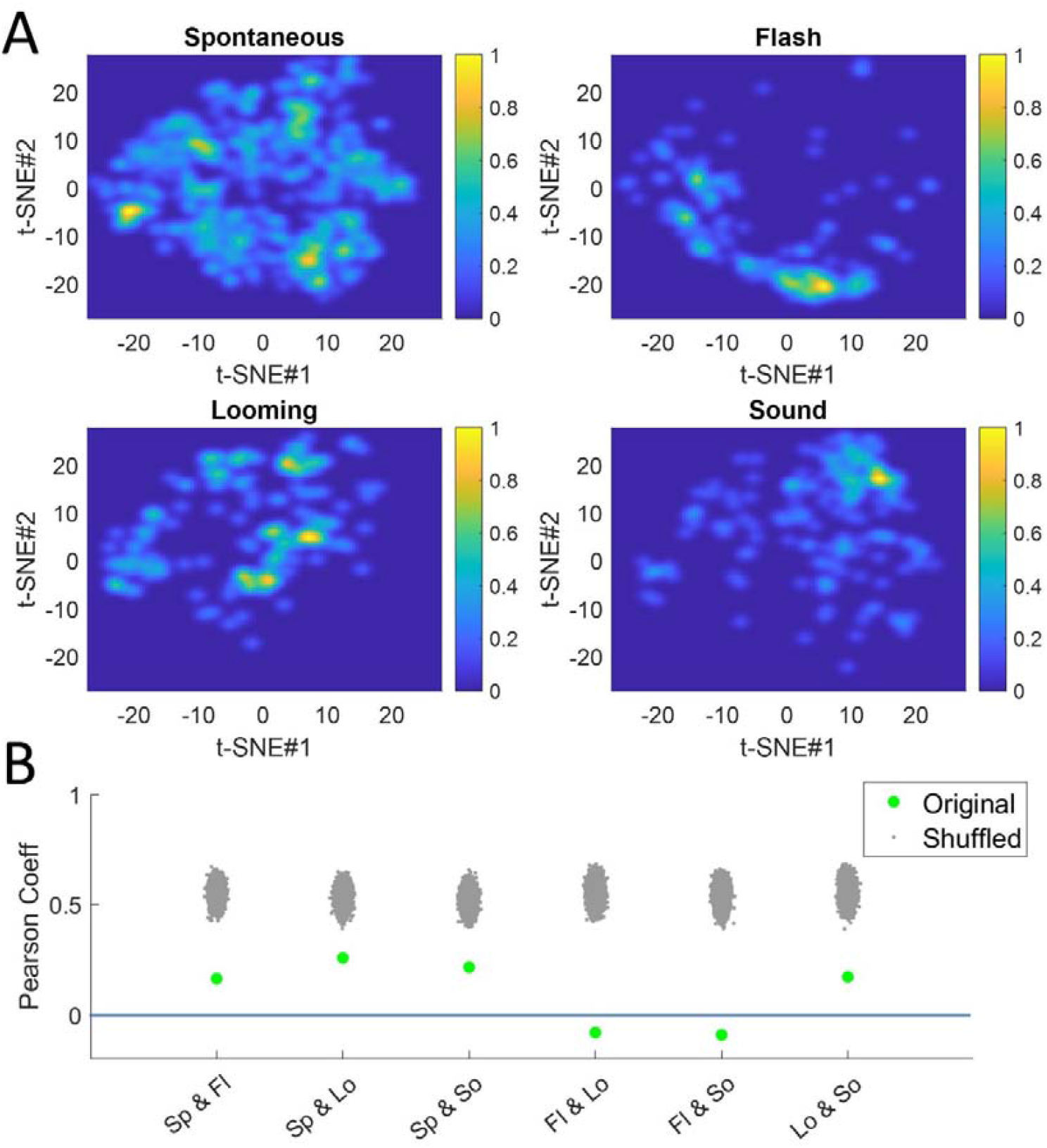
**A)** Each panel shows map representation of different behavioural states. Maps are obtained by calculating a 2D count histogram of a 2-dimensional t-SNE embedding shown in **Figure 4C**. The different maps represent spontaneous exploration (top left) and the three different stimulation conditions (Flash, Looming, Sound). X and Y axes here represent the first and second t-SNE embedding dimensions (respectively t-SNE#1 and t-SNE#2). The maps values, ranging from 0 to 1, are obtained by normalising each histogram by its highest count. **B)** Overlap between pairs of maps are quantified as Pearson’s correlation between maps (green points; Sp = Spontaneous; Fl = Flash; Lo = Looming; So = Sound). Across all pairs the amount of overlap is substantially smaller than expected by chance since shuffling stimulus conditions returns larger correlations values (grey points, 10000 shuffles for each pair).

To quantitatively test these initial observations, we performed a set of statistical shuffle tests across all pairs of t-SNE maps. We first quantified the overlap between two t-SNE maps by calculating their Pearson’s correlation coefficient (**Figure 4B**, green dots). We then randomly shuffled the t-SNE points across pairs of maps to determine the level of chance overlap. Shuffling was performed by using individual animals as blocking factor to avoid introducing additional interindividual variability. We found that the overlap measured on the shuffled data was always larger than that measured on the original data (**Figure 4B**, grey dots; shuffle repeats = 10000, p-value = 0 across all pairs of t-SNE maps). These results indicate that behavioural responses to different stimuli are strongly segregated from each other and from spontaneous behaviours.

## Discussion

Reduction and Refinements two of the 3Rs principles outlined to provide a framework for more humane animal research. We described here how to build and set-up a system for 3D reconstruction of mouse freely moving behaviour. This enables detailed and reproducible quantification of freely moving animal behaviours a key requirement for reducing animal numbers and avoid more severe procedures based on head fixations and food/water deprivation. Studying natural unconstrained behaviours has also fundamental implications for advancing neuroscience. The need to break away from reductionist approaches to progress fundamental research has been acknowledged over the last few years [31-33].

To address these welfare and scientific problems, over the last few years other systems have also been developed to perform 3D reconstructions of freely moving animals across different species (e.g. rodents [24, 34], monkeys [35] and cheetahs [4]). Here we provide a detailed description of the material and procedures to build and calibrate such a 3D system for mice. We hope the information included here will help other groups to adopt 3D technologies for behavioural studies in rodents.

## Notes

### Competing Interest Statement

The authors have declared no competing interest.

